# Chronic and Acute Mediators of Passive Viscoelasticity in Human Skeletal Muscle Fibers

**DOI:** 10.1101/2024.08.13.607865

**Authors:** Grace E. Privett, Austin W. Ricci, Karen Wiedenfeld Needham, Damien M. Callahan

**Affiliations:** Department of Human Physiology, University of Oregon, Eugene, OR, USA

**Keywords:** skeletal muscle, fatigue, training, stress decay, viscoelasticity, passive mechanics, cellular stiffness, titin

## Abstract

Cellular viscoelastic modulus in skeletal muscle tissue responds dynamically to chronic stressors, such as age and exercise. Passive tissue mechanics may also be sensitive to acute stimuli such as mechanical loading and/or activation-induced muscle fatigue. These insights are largely derived from preclinical studies of age and acute muscle activation. Therefore, we sought to understand the relative responsiveness of muscle cellular passive mechanics to chronic (resistance training) and acute (muscle fatigue) stressors in healthy young males and females categorized as “resistance trained” or “untrained”. We measured passive mechanics to test the hypothesis that Young’s Modulus and stress would be greater in fibers from trained versus untrained participants and both would be reduced following fatigue. We further assessed the translation of these findings to composite tissue in a sub-set of volunteers where muscle tissue bundles, containing both fibers and extracellular matrix, were analyzed in addition to single fibers. We report a main effect of training such that cellular passive mechanical measures were increased in single fibers from trained versus untrained participants. We likewise report reductions in passive mechanical measures following fatiguing exercise. Surprisingly, both training and acute fatigue only impacted muscle fiber passive measures in males, whereas females showed a more variable response across conditions. Last, we provide preliminary evidence supporting the translation of per-individual cellular differences to the tissue level. Together, these data suggest males respond more dynamically to acute and chronic stressors of muscle tissue mechanics, potentially linking cellular response and sex-dependent differences in musculotendinous injury risk.

**New and noteworthy:** We report that passive stress and modulus in single muscle fibers was higher in resistance trained healthy adults and fatiguing exercise reduced passive stress and modulus. In each case, dynamic responsiveness of muscle fibers to chronic and acute stressors was observed consistently in males, whereas responses in females varied considerably. We provide further evidence that cellular mechanisms may contribute to multicellular muscle tissue samples, suggesting these findings have relevance to in vivo tissue mechanics.

## Introduction

Skeletal muscle viscoelasticity affects functional mobility through multiple interrelated mechanisms including its influence on joint stability (Blackburn *et al*., 2011), and the dampening of impact forces (Sarvazyan *et al*., 2014) during musculoskeletal loading. Viscoelastic behavior of muscle also contributes to the rate and mechanical efficiency of force transfer from muscle to tendon (Wilson & Flanagan, 2008). Prior studies report that whole-muscle stiffness and viscosity are transiently altered by a single bout of fatiguing exercise in the Vastus Lateralis (VL) (Chalchat *et al*., 2020). This transient change to VL mechanical properties may be maladaptive, considering that muscular fatigue was previously shown to decrease the capacity for skeletal muscle to absorb strain elastic energy before incurring injury (Mair *et al*., 1996). As such, acute, fatigue-induced changes to skeletal muscle viscoelasticity may contribute to the high prevalence of hamstring injury in male athletes (Watsford *et al*., 2010; Cross *et al*., 2013) or knee joint destabilization (Blackburn *et al*., 2011) and subsequent ACL injury in female athletes (Myer *et al*., 2008; De Ste Croix *et al*., 2015). Given the likely relationship between muscle viscoelastic properties and tissue injury risk, it is logically appealing to predict that biological sex, muscle fatigue and injury risk share a complex dynamic contributing to sex-disparity in rates of soft tissue injury.

At the cellular level, we have demonstrated fatigue-induced reductions in passive Young’s Modulus, a measure of stiffness accounting for differences in cell (“fiber”) size, in VL samples from young, recreationally active adults in a sex-dependent manner (Privett *et al*., 2024). At the single fiber level, passive modulus is largely attributed to the intracellular protein titin (Ottenheijm *et al*., 2012; Lim *et al*., 2019). Titin is a viscoelastic protein that is subject to post-translational modifications (PTM), many of which can alter titin-based stiffness if they occur at an extensible region of the protein (Hamdani *et al*., 2017). Ample study of PTM-induced changes to titin-based mechanical properties has been conducted in cardiac muscle, which has demonstrated that phosphorylation of cardiac titin regulates cardiomyocyte mechanics in health and disease (Granzier & Irving, 1995; Wu *et al*., 2000, 2002; Yamasaki *et al*., 2002; Fukuda *et al*., 2005; Krüger *et al*., 2009; LeWinter & Granzier, 2014). Skeletal muscle titin can also be phosphorylated (Müller *et al*., 2014; Privett *et al*., 2024), though whether titin phosphorylation alters viscoelasticity in skeletal muscle is not yet clear. Exercise is a multimodal and dynamic physiological stimulus that likely induces multiple PTMs to consider in skeletal muscle. These include phosphorylation (Müller *et al*., 2014), S-glutathionylation (Alegre-Cebollada *et al*., 2014; Watanabe *et al*., 2020) and the binding of small heat shock proteins (HSP, Kötter *et al*., 2014), the last of which has been shown to protect against aggregation of titin immunoglobulin domains and subsequent stiffening of the protein (Kötter *et al*., 2014). Evidence suggests that titin directly influences whole muscle function (Brynnel *et al*., 2018), supporting the clinically relevant translation of our recent observation that acute regulation of titin may influence skeletal muscle mechanics (Privett *et al*., 2024). Therefore, the current study sought to test the role of intracellular contributors to changes in tissue-level mechanical properties by comparing the impact of acute fatigue on viscoelastic properties of single fibers (titin-based) and in bundles of fibers with intact extracellular matrix (ECM) who’s modulus is presumably impacted by several factors outside the myofiber.

Although the study of altered VL mechanical properties following fatiguing exercise may have high clinical relevance for athletes, it remains to be seen whether the neuromuscular adaptations to chronic training mediate these observations. Chronic training has been demonstrated to increase active (Mongold *et al*., 2022) and resting (Klinge *et al*., 1997) musculoskeletal stiffness in some, but not all (Blazevich, 2019), studies. Evidence of training-based changes to collagen expression, synthesis, and accumulation in the ECM (Kjær, 2004) suggests that altered skeletal muscle stiffness due to training may arise from modified ECM-based stiffness. Although it is unclear whether chronic training impacts ECM-based measures of stiffness in the literature, jump training in rats has been shown to increase the passive stiffness of extensor digitorum longus (EDL) and rectus femoris (RF) muscles in conjunction with increased collagen concentration (Ducomps *et al*., 2003). At the cellular level, training was shown to alter the stiffness of permeabilized skeletal muscle fibers in a length-dependent manner (Noonan *et al*., 2020) suggesting that intracellular mechanisms may also contribute to training-based changes in skeletal muscle stiffness. Therefore, the present study compared baseline measures of cellular passive viscoelasticity in VL samples from untrained (UT) and resistance trained (RT) young males and females and expanded upon our previous work (Privett *et al*., 2024) by assessing whether chronic training mediates the effect of fatiguing exercise on cellular passive mechanics.

Although the majority of previous research focused on measures of elasticity, viscosity is equally important to consider given its role in skeletal muscle extensibility and the absorption and dissipation of external mechanical loads (Sarvazyan *et al*., 2014). Therefore, this present study sought to extend our previous work by considering the effect of fatigue on the viscoelasticity of human skeletal muscle cells. To do so, stress decay index (SDI) was calculated using the magnitude and rate of stress relaxation at each sarcomere length (SL) studied.

The purpose of this study was to compare passive viscoelastic properties in VL samples obtained from exercised (“fatigued”) and non-exercised (“non-fatigued”) limbs of UT and RT young adults. We hypothesized that passive mechanical measures (stress, Young’s Modulus) would be reduced in fatigued versus non-fatigued skeletal muscle fibers from all participants. Based on evidence of training-induced stiffness increases in the literature (Noonan *et al*., 2020), it was also hypothesized that cellular passive mechanical measures would be higher in RT versus UT individuals. To consider the “viscous” element of skeletal muscle viscoelasticity, SDI was compared in fatigued and non-fatigued skeletal muscle cells.

Finally, passive modulus was measured in bundles of fibers with intact ECM, termed here as “composite muscle tissue” (CMT) before and after chemical elimination of intracellular contributors to passive modulus to assess the extent to which intracellular mechanisms contribute to observations of altered CMT passive modulus after fatiguing exercise. It was hypothesized that any effect of fatiguing exercise on measures of CMT stiffness would be abolished by elimination of intracellular contributors to passive modulus.

## Methods

### Population

This protocol was approved by the Institutional Review Board at the University of Oregon. 19 young (aged 21 ± 3 yrs.), RT and UT males and females from the University of Oregon and surrounding community consented to participate in this study. Participants were considered RT if they reported between five and seven sessions, each at least 1-hour in duration, of weight training, with at least three sessions focused on the lower limbs. Participants were considered UT if they reported recreational levels of activity with no participation in structured physical exercise and no resistance training. Initial self-report of physical activity history was confirmed by use of ActivePal accelerometers (Glasgow, Scotland) as described previously (Dowd *et al*., 2012). To limit the potential for menstrual cycle-dependent variation in circulating estradiol and associated potential impacts on skeletal muscle mechanical properties, all female volunteers either reported use of hormonal contraceptive or were tested in the pre-follicular phase of the menstrual cycle, (within 5 days of menses onset). Participants reported no orthopedic limitations (severe osteoarthritis, joint replacement, or other orthopedic surgery in the previous six months), endocrine disease (hypo/hyper thyroidism, Addison’s Disease or Cushing’s syndrome), uncontrolled hypertension (>140/90 mmHg), neuromuscular disorder, significant heart, liver, kidney or respiratory disease, or diabetes. Participants were non-tobacco-smokers and had no current alcohol disorder. Finally, participants taking medications known to affect muscle stiffness or beta-adrenergic signaling of neuromuscular activation (including but not limited to beta blockers, calcium channel blockers, and muscle relaxers) or anabolic steroids were not included.

### Study Design

Participants visited the lab on 2 occasions separated by at least 1 week. During the first visit, non-invasive measures of voluntary strength, power, and fatigue of their dominant knee extensors (KE) were collected using a Biodex System 3 dynamometer (Biodex Medical Systems, Shirley, NY) and participants were familiarized with the exercise protocol. During the second visit, volunteers performed maximal voluntary isometric contractions of the KE followed by fatiguing exercise to task failure. Fatiguing exercise was followed by bilateral, percutaneous needle muscle biopsies: one on the exercised limb immediately following exercise (“fatigued”) and the second on the contralateral, non-exercised limb (“non-fatigued”).

### Fatigue Protocol

The fatigue protocol utilized for this study was described in detail previously (Privett *et al*., 2024). Briefly, Participants were seated on the Biodex dynamometer with hips and knee flexed at 90° (180° = full extension). After completion of three maximum voluntary isometric contractions (MVIC) of the knee extensors of the dominant leg, participants performed repeated, voluntary knee extensions to task failure while the Biodex applied a load set to 30% MVIC maximal torque. Task failure was identified as the inability to perform knee extension through at least 50% range of motion. Fatigue was quantified as the Fatigue Ratio 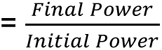, where “initial power” represents the average peak power of the first five knee extensions performed during fatiguing exercise, and “final power” represents the average peak power from the last five knee extensions. Time to fatigue (task failure) was recorded for all participants.

### Muscle Biopsy Procedure

Percutaneous needle biopsy of the VL was performed within 9 ± 4 minutes following task failure, under sterile conditions as previously detailed (Tarnopolsky *et al*., 2011). First, the biopsy site was sterilized and local anesthetic (1 or 2% lidocaine HCL [Hospira Worldwide, Lake Forest, IL, USA]) was administered via injection. Next, a small (∼5 mm) incision was made in the skin and muscle fascia, allowing for the passage of a Bergstrom biopsy needle (5 mm diameter) to the belly of the VL muscle to acquire sample at a depth of ∼2-3 cm. Following acquisition of biopsy sample, the collected muscle was retrieved from the needle using forceps.

### Tissue Processing and Dissection

The details of tissue processing for mechanics were detailed elsewhere (Privett *et al*., 2024). Briefly, sample collected during biopsy was placed in muscle dissecting solution (MDS, 120.782 mM sodium methanesulfonate (NaMS), 5.00 mM EGTA, 0.118 mM CaCl_2_, 1.00 mM MgCl_2_, 5.00 mM ATP-Na_2_H_2_, 0.25 mM KH_2_PO_4_, 20.00 mM BES, 1.789 mM KOH, 1 mM Dithiothreitol (DTT)), parsed into bundles of ∼50 fibers, and tied to glass rods before advancement through solutions of increasing glycerol content and storage in 50% glycerol solution (5.00 mM EGTA, 2.50 mM MgCl_2_, 2.50 mM ATP-Na_2_H_2_, 10 mM imidazole, 170.00 mM potassium propionate, 1.00 mM sodium azide, 50% glycerol by volume) at -20°C. Sample allocated for mechanics analyses were used within 4 weeks following biopsy. Preparation for mechanical assays was described previously (Privett *et al*., 2024).

Briefly, fiber bundles and dissected single fibers were chemically skinned (MDS + 1% Triton X-100), transferred to plain MDS, and kept on ice until experimentation. For CMT experiments, fiber bundles were transferred directly from 50% glycerol solution to plain MDS (no further chemical skinning). Then, strips of 12-14 fibers with surrounding ECM were dissected from the bundle and kept on ice until experimentation.

### Single fiber morphology and contractile measures

Prepared fibers were mounted in relaxing solution (67.286 mM NaMS, 5.00 mM EGTA, 0.118 mM CaCl_2_, 6.867 mM MgCl_2_, 0.25 mM KH_2_PO_4_, 20.00 mM BES, 0.262 mM KOH, 1.00 mM DTT, 5.392 mM Mg-ATP, 15.00 mM creatine phosphate (CP), 300 U/mL creatine phosphokinase (CPK)) between a force transducer and a length motor (Aurora Scientific, Inc., Aurora, ON, Canada) using the Moss clamp technique (Moss, 1979). Passive tension was measured by zeroing the force transducer while the fiber was slacked, then stretching the fiber to SL = 2.65 µm.

Active tension was measured at SL 2.65 µm by moving the fiber to pre-activating solution (81.181 mM NaMS, 5.00 mM EGTA, 0.012 mM CaCl_2_, 6.724 mM MgCl_2_, 5.00 mM KH_2_PO_4_, 20.00 mM BES, 1.00 mM DTT, 5.397 mM Mg-ATP, 15.00 mM CP, 300 U/mL CPK) followed by activating solution (57.549 mM NaMS, 5.00 mM EGTA, 5.021 mM CaCl_2_, 6.711 mM MgCl_2_, 5.00 mM KH_2_PO_4_, 20.00 mM BES, 9.674 mM KOH, 1.00 mM DTT, 5.437 mM Mg-ATP, 15.00 mM CP, 300 U/mL CPK). Once a steady state tension was recorded, the sample was returned to relaxing solution. All fibers were activated (pCa 4.5) prior to passive stretching to measure active tension and confirm fiber viability.

### Passive stretch protocol

Passive stretch measures were performed in relaxing solution (pCa 8.0) using a passive stretch protocol (Figure 1) adapted from previous work (Lim *et al*., 2019). Initial sarcomere length was set to 2.4 µm, followed by 7 incremental stretches to reach a final length of 156% of initial length (SL = 3.5-4.0 µm). Each stretch lengthened the sample 8% of initial length and held this position for 2 minutes of stress-relaxation. SL was measured throughout the protocol using an inverted microscope located beneath the single fiber rig. To determine the extent to which actomyosin interactions contributed to measures of passive viscoelasticity, a subset of fibers from two sedentary males was subjected to the passive stretch protocol in relaxing solution with the addition of 40 mM 2,3-butanedione monoxime (BDM), a myosin inhibitor. Following completion of the passive stretch protocol, each fiber was collected and placed in gel loading buffer (2% SDS, 62.5 mM Tris, 10% glycerol, 0.001% bromophenol blue, 5% β-mercaptoethanol, pH 6.8), centrifuged and heated at 65°C for 2 minutes, then stored at -80°C until later assessment of myosin heavy chain (MHC) isoform.

**Figure 1:**
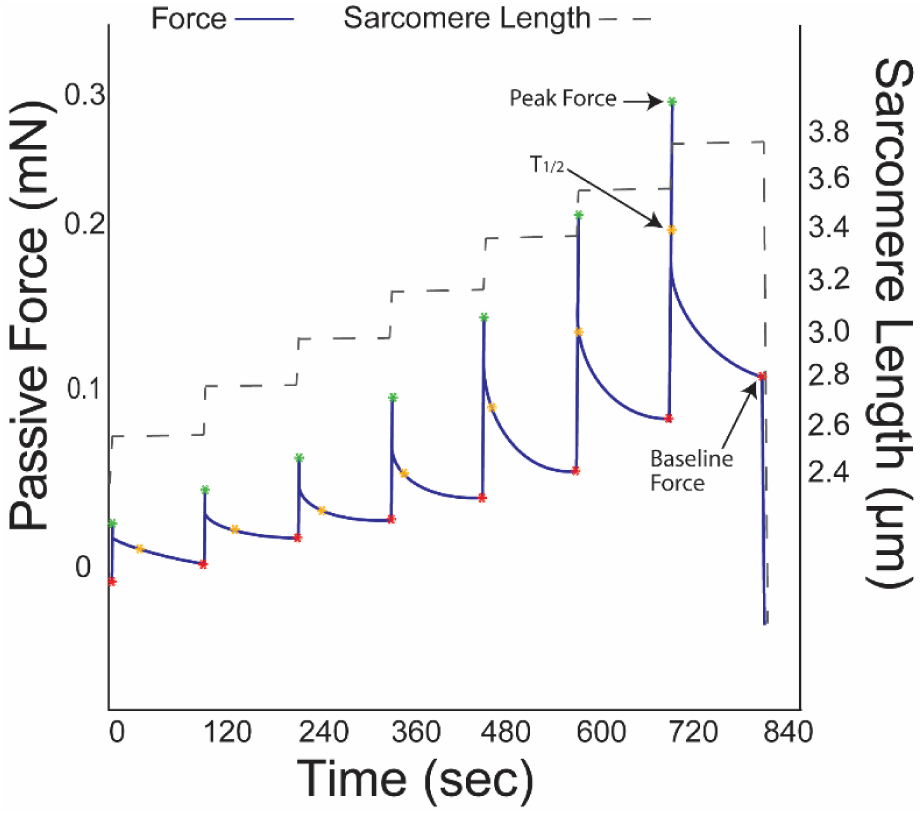
Sample force trace indicating the outcome measures produced by this passive stretch protocol. Baseline force was used to calculate passive stress and peak force, half-relaxation time (T_1/2_), and baseline force were used to calculate SDI.

*KI/KCl treatment*: To assess titin’s contribution to passive modulus in CMT, mounted bundles of ∼12-14 fibers with intact ECM were treated with potassium chloride (KCl) and potassium iodide (KI) to extract the thick and thin sarcomeric filaments, respectively (Ottenheijm *et al*., 2012; Brynnel *et al*., 2018). CMT samples were first subjected to an initial passive-stretch (as described above), after which the sample was incubated in relaxing solution containing 0.6 M KCl for 35 minutes at 15°C followed by relaxing solution containing 1.0 M KI for 35 minutes at 15°C. After incubation, CMT samples were passively stretched a second time to collect post-treatment passive mechanical measures. Following these experiments, CMT samples were collected and stored in gel loading buffer.

### MHC isoform identification

Sodium dodecyl sulfate poly acrylamide gel electrophoresis (SDS-PAGE) was used to determine the MHC isoform of single muscle fibers. Sample from each fiber was loaded into its own well of a 4% stacking / 7% resolving polyacrylamide gel. The gel was run at 70 V for 3.5 hours followed by 200 V for 20 hours at 4°C (Miller *et al*., 2010). Gels were stained with silver and the resulting MHC isoform (I, IIA, and/or IIX) expression was determined by comparison to a standard made from a multi-fiber homogenate (Figure 2).

**Figure 2:**
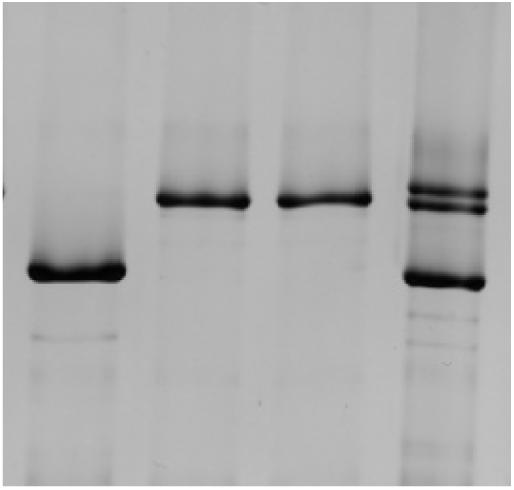
Sample image of silver-stained MHC bands. The left lane shows a fiber exhibiting MHC I isoform, the two middle bands exhibit MHC IIA isoform, and right lane contains a sample homogenate used to visualize all three (I, IIA, IIX) isoforms.

### Outcome Measures

Maximally activated tension was quantified as the plateau of active force divided by fiber CSA. Passive stress at each sarcomere length (SL) was calculated as 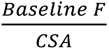, where “Baseline F” indicates force following stress relaxation (Figure 1) and “CSA” indicates fiber cross sectional area assuming elliptical shape. 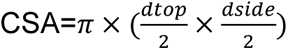, where “d_top_” is the average of three top diameter measures, and “d_side_” is the average of three side diameter measures. Strain was calculated as 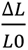, where “L_0_” indicates initial length. Using raw stress data, passive Young’s Modulus was calculated as the slope of the stress-strain relationship at shorter fiber lengths (strain = 1.0-1.24 %L_0_) and at longer fiber lengths (strain = 1.32-1.56 %L_0_). For fiber measures of passive modulus, separate slopes were calculated for shorter and longer fiber lengths to consider the length dependence of passive modulus measures. Stress decay index (SDI) was calculated as previously described (Lim *et al*., 2019): 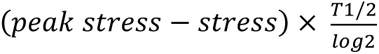. 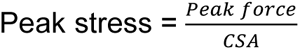 and T_1/2_ = half-relaxation time (Figure 1). For tissue-level measures, passive Young’s Modulus was calculated as one slope of the stress-strain curve, given the lack of a clear curvilinear inflection point such as that observed in the stress-strain data of the fibers. The direction and magnitude of change in modulus between non-fatigued and fatigued samples was calculated for fiber and CMT samples as 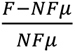, where “F” indicates the fatigued value for each individual sample and “NF_µ_” indicates the group mean of the non-fatigued samples. For KI/KCl passive measures of CMT, passive modulus was calculated as one slope of the stress-strain curve.

### Statistical Analyses

Statistical testing was conducted using SPSS software package (SPSS, IBM Corp., Armonk, NY, USA) unless otherwise specified. At each SL, differences in passive stress and SDI were evaluated using separate linear mixed models including fatigue, biological sex, training, and interaction terms as fixed effects and participant ID as a random effect to account for fiber variation within individuals, as described previously (Callahan *et al*., 2015). To evaluate differences in single fiber passive modulus at short and long lengths, separate linear mixed models were run with fatigue, biological sex, training and interaction terms as fixed effects and participant ID as a random effect. To test for differences in maximally activated tension, a mixed effects model was run including sex, training, and fatigue as main effects and participant ID as a random effect. Of note, because MHC IIA fibers were similarly represented across all groups, all statistical analyses for single fiber mechanics included only MHC IIA fibers. Though this decision precluded the consideration of interactions between fiber type and other main effects (sex/fatigue) in our cohort (See Table 2), we considered this to be the most rigorous and conservative approach to test our hypotheses with respect to sex and fatigue, though we acknowledge it limits fiber number somewhat (206 fibers out of 423, total). To test for significant differences in the passive Young’s Modulus of CMT samples, a linear mixed model was generated with biological sex and fatigue as fixed effects and participant as a random effect. Follow-up analyses tested for an effect of fatigue, with participants as a random effect, on passive modulus of CMT samples from males and females, separately. To test for fatigue-based differences in CMT passive modulus before and after KI/KCl treatment, means for the 4 fatigue/treatment groups (non-fatigued pre-KI/KCl, non-fatigued post-KI/KCl, fatigued pre-KI/KCl, fatigued post-KI/KCl) were compared in a univariate analysis of variance with least significance difference (LSD) post-hoc testing.

## Results

### Descriptive Measures

There were significant main effects of biological sex on height and weight such that males were taller than females (184.3 ± 8.6 vs. 161.6 ± 4.9 cm, respectively, p<0.01) and weighed more than females (81.3 ± 9.3 vs. 57.4 ± 5.2 kg, respectively, p<0.01, Table 1). There was no significant main effect of training on height (p=0.691) or weight (p=0.997). BMI was significantly higher in males versus females (23.9 ± 1.7 versus 22.0 ± 1.8 kg/m^2^, respectively, p=0.023), but was not different between RT and UT participants (p=0.463). Regarding activity levels, RT participants exhibited higher step count (12505 ± 2823 vs. 7201 ± 2162 steps, respectively, p<0.01) and minutes in moderate physical activity (101.8 ± 19.3 vs. 54.0 ± 16.0 minutes per day, respectively, p<0.01). There was no effect of biological sex on step count (p=0.928) or time spent in moderate physical activity (p=0.614). There was no significant main effect of training status or biological sex on minutes in light physical activity (p=0.090 and 0.185, respectively) or minutes in vigorous physical activity (p=0.183 and p=0.400, respectively). In total, data were collected and analyzed for 424 fibers (Table 2). Single fiber CSA was greater in males versus females (0.0074 ± 0.0024 vs. 0.0050 ± 0.0016 mm^2^, respectively, p<0.01) and greater in RT versus UT participants (0.0065 ± 0.0025 vs. 0.0055 ± 0.0019 mm^2^, respectively, p=0.047). There was no effect of biological sex (p=0.344) or training (p=0.484) on fiber length. MHC isoform of individual fibers was determined using SDS-PAGE (Figure 2). Fiber type distribution differed between males and females such that females exhibited a greater proportion of MHC I fibers than males (28% versus 6%, respectively). In both groups, MHC II (including IIA, IIX, and A/X) fibers comprised the majority of the sample (72% in females, 94% in males). Because MHC IIA fibers were similarly represented across all groups, all statistical analyses for single fiber mechanics included only MHC IIA fibers, as mentioned above.

**Table 1:** Anthropometric and Activity Data of Included Participants.

|  |  | n | Height (cm) * | Weight (kg) * | BMI (kg/m<sup>2</sup>) * | Step Count (steps/day) # | Light Activity (mins/day) | Moderate Activity (mins/day) # | Vigorous Activity (mins/day) |
| --- | --- | --- | --- | --- | --- | --- | --- | --- | --- |
| UT | Female | 5 | 162.1 ± 6.1 | 55.0 ± 3.2 | 20.9 ± 0.6 | 7274±2162 | 27 ± 9 | 56 ± 18 | 0.5 ± 0.6 |
|  | Male | 4 | 185.3 ± 13.0 | 84.0 ± 10.1 | 24.4 ± 1.2 | 7108±1525 | 38 ± 7 | 51 ± 15 | 1.1 ± 1.2 |
| RT | Female | 5 | 160.1 ± 3.9 | 60.0 ± 6.0 | 23.1 ± 2.1 | 12357±2531 | 40 ± 13 | 96 ± 16 | 5.0 ± 6.2 |
|  | Male | 5 | 183.6 ± 4.5 | 79.1 ± 9.2 | 23.4 ± 2.1 | 12752±3864 | 43 ± 12 | 111 ± 24 | 1.5 ± 1.3 |

Symbols indicate a significant main effect of \* biological sex or # training between groups (p<0.05). Data are shown as mean ± SD.

**Table 2:** Summary Statistics of Fibers Analyzed.

|  |  | n | CSA (mm<sup>2</sup>)<br>*, # | Length<br>(mm) | Relative % MHC |  |  |  |
| --- | --- | --- | --- | --- | --- | --- | --- | --- |
|  |  |  |  |  | I | IIA | IIX | IIA/X |
| UT | Female | 107 | 0.0046 ± 0.0014 | 1.5 ± 0.4 | 22 | 47 | 2 | 22 |
|  | Male | 92 | 0.0066 ± 0.0019 | 1.7 ± 0.4 | 3 | 49 | 23 | 24 |
| RT | Female | 128 | 0.0053 ± 0.0016 | 1.6 ± 0.6 | 34 | 47 | 2 | 12 |
|  | Male | 97 | 0.0081 ± 0.0026 | 1.8 ± 0.5 | 9 | 55 | 2 | 34 |

Data are shown as mean ± SD. Percents are not reported for MHC I/IIA or I/IIA/IIX because they collectively made up only 3.1% of the overall dataset. Symbols indicate main effects of \*biological sex or # training (all p<0.05).

### Fatiguing exercise and whole-muscle contractile measures

Due to technical limitations related to data loss during transfer, time to fatigue and knee extensor power measures are reported for 18/19 and and 16/19 volunteers, respectively. The average time to fatigue was not significantly different by training status (p=0.453) or biological sex (p=0.559). Similarly, there was no effect of training status (p=0.415) or biological sex (0.926) on fatigue ratio (Table 3). Absolute peak power was significantly higher in RT versus UT (608.35 ± 295.87 versus 413.84 ± 143.11 W, respectively, p=0.017) and male versus female (683.13 ± 238.86 versus 339.06 ± 70.27 W, respectively, p<0.001) participants. Relative peak power was significantly higher in RT versus UT (8.27 ± 2.58 versus 5.93 ± 1.09 W/kg, respectively, p=0.002) and male versus female (8.40 ± 2.60 versus 5.81 ± 0.64 W, respectively, p=0.001) participants. Absolute peak torque was significantly higher in RT versus UT (279.55 ± 102.78 versus 202.45 ± 79.51 N, respectively, p=0.004) and male versus female (327.32 ± 60.12 versus 167.16 ± 50.97 N, respectively, p<0.001) participants. Finally, relative peak torque was significantly higher in RT versus UT (3.91 ± 0.96 versus 2.90 ± 0.48 N/kg, respectively, p=0.001) and male versus female (4.06 ± 0.80 versus 2.87 ± 0.60 N/kg, respectively, p<0.001) participants.

**Table 3:** Fatigue data and whole-muscle performance.

|  |  | Time to<br>Fatigue (sec) | Fatigue<br>Ratio | Peak Power<br>(W) ** | Relative Peak<br>Power (W/kg) ** | Peak Torque<br>(N) ** | Relative Peak<br>Torque (N/kg) ** |
| --- | --- | --- | --- | --- | --- | --- | --- |
| UT | Female | 68±8.8 | 0.34±0.10 | 295±33 | 5.43±0.32 | 140±8 | 2.54±0.12 |
|  | Male | 75±29 | 0.41±0.16 | 532±95 | 6.43±1.42 | 281±44 | 3.34±0.34 |
| RT | Female | 99±74 | 0.36±0.12 | 383±73 | 6.19±0.68 | 195±62 | 3.20±0.73 |
|  | Male | 71±14 | 0.28±0.10 | 834±252 | 10.37±1.85 | 364±44 | 4.62±0.53 |

Symbols indicate a significant main effect of \*biological sex or #training between groups (p<0.05). Data are shown as mean ± SD.

### Maximal Isometric Tension

Considering the known impact of MHC isoform on active tension, and the relative balance in proportion of MHC IIA fibers across groups, maximally active isometric tension was analyzed in MHC IIA fibers only. There was no main effect of training (p=0.219), biological sex (p=0.843), or fatigue condition (p=0.653) on maximal isometric tension; however, the interaction between biological sex and fatigue was significant (p=0.013) such that maximal tension was reduced modestly (∼5%) by fatiguing exercise in males (174.0 ± 37.7 vs. 164.7 ± 46.2 kPa, respectively p=0.021) but not females (178.1 ± 36.4 vs. 179.0 ± 37.1 kPa, respectively, p=0.163, Figure 3).

**Figure 3:**
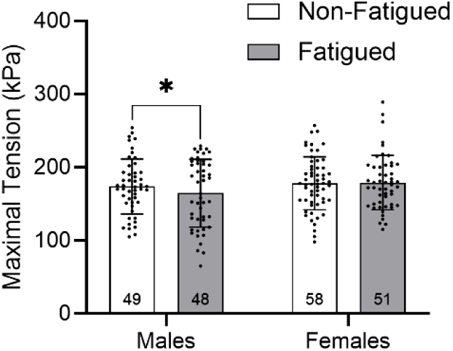
Active tension generation was assessed in MHC IIA fibers. There was no significant effect of training, biological sex, or fatigue condition on active tension. Active tension was significantly reduced in fatigued fibers of males but not females. Data are shown as mean ± SD.

### Assessment of Passive Stretch Protocol

To address potential concerns regarding incomplete stress decay after each applied strain, the difference between the measured stress and predicted stress, termed “missed stress”, was calculated at each stretch step, for each fiber as described previously (Privett *et al*., 2024). Stress decay was greatest at SL=3.2-3.8 µm, suggesting the greatest potential for incomplete stress decay. At these lengths, measured stress was only different from predicted stress by 0.08-2.37%. Given this minimal variation, it is not likely that incomplete stress decay impacted the ability to draw meaningful conclusions from final measured stress, thus measured stress was used throughout.

*Passive Stress:* Passive stress was analyzed at each step of the passive stretch protocol. When all MHC IIA fibers were considered (Figure 4A), passive stress was significantly different by fatigue (p=0.034 at SL=2.6 µm, p<0.01 at SL=2.8-3.8 µm), fatigue by training interactions (p= 0.049 at SL 2.6 µm, p= 0.038 at SL 2.8 µm, p= 0.028 at SL 3.0 µm, p= 0.023 at SL 3.2 µm, p= 0.017 at SL 3.4 µm, p= 0.013 at SL 3.6 µm, p=0.017 at SL 3.8 µm), and fatigue by sex interactions (p<0.01 at all SL). When considering only non-fatigued fibers (Figure 4B), passive stress was significantly higher in RT versus UT fibers at longer SL (p= 0.033 at SL 3.0 µm, p= 0.035 at SL 3.2 µm, p= 0.034 at SL 3.4 µm, p= 0.032 at SL 3.6 µm, p=0.034 at SL 3.8 µm). There was no main effect of biological sex on passive stress in non-fatigued fibers, at any SL. In UT males (Figure 4C), passive stress was significantly lower in fatigued versus non-fatigued fibers at longer lengths (p= 0.032 at SL 3.0 µm, p= 0.018 at SL 3.2 µm, p= 0.018 at SL 3.4 µm, p= 0.010 at SL 3.6 µm, p<0.01 at SL 3.8 µm). In UT females (Figure 4D), passive stress was not significantly different by fatigue at any SL. Furthermore, in fibers from UT participants, the effect of fatiguing exercise on passive stress differed by biological sex at longer SL (p= 0.019 at SL 3.0 µm, p<0.01 at SL 3.2-3.8 µm). In RT males (Figure 4E), passive stress was significantly reduced by fatigue at all SL (p<0.01 at SL 2.6-3.8 µm) and the response of passive stress to fatigue was greater than that of UT males (interaction p= 0.018 at SL 2.6 µm, p<0.01 at SL 2.8-3.0 µm, p= 0.010 at SL 3.2 µm, p<0.01 at SL 3.4-3.8 µm). In fibers from RT females (Figure 4F), fatigue did not significantly affect passive stress at any SL, and the response of passive stress to fatiguing exercise was different from that of RT males at all SL (p<0.01 at SL 2.6-3.8 µm).

**Figure 4:**
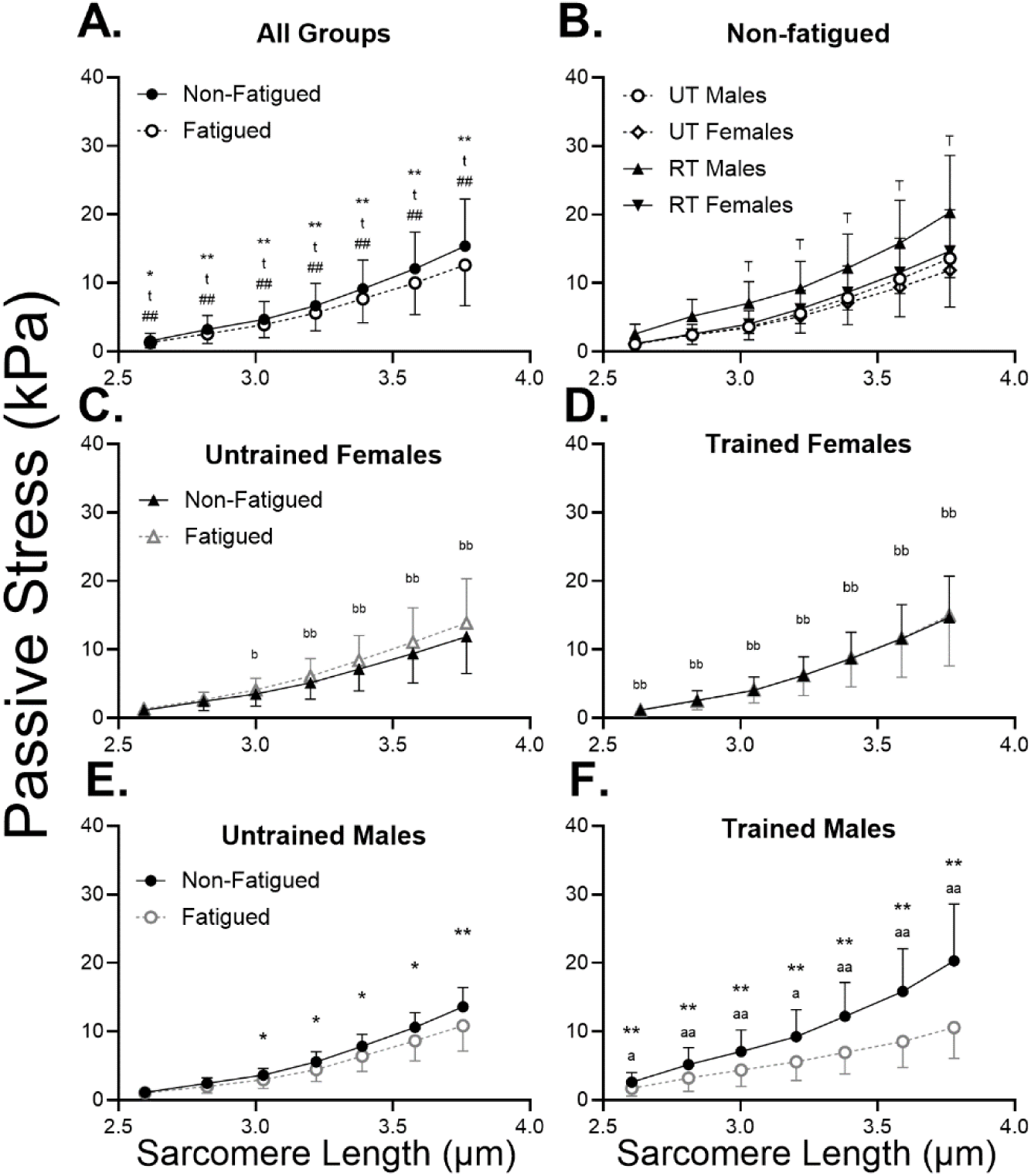
In the combined group, passive stress was significantly reduced at all sarcomere lengths, though the response was different by biological sex and training group **(A)** In non-fatigued fibers, passive stress was significantly higher in RT versus UT participants **(B)**. In UT participants, passive stress was significantly reduced by fatiguing exercise at SL=3.0-3.8 µm in males **(C)** but not females **(D)**, due to significant fatigue by sex interactions at these lengths. In RT participants, the effect of fatigue on passive stress was highly significant at all SL in males **(E)** but not in females **(F)**, due to the highly significant fatigue by sex interaction in this group. Furthermore, the response of passive stress to fatiguing exercise was different between RT and UT males. Data are shown as mean ± SD. Symbols indicate a significant effect of fatigue (* p<0.05 or ** p<0.01), fatigue by training interaction (^t^ p<0.05), fatigue by biological sex interaction (^#^ p<0.05 or ^##^p<0.01), main effect of training (^T^ p<0.05), fatigue response different from that of UT (^a^ p<0.05 or ^aa^ p<0.01), fatigue response different from that of males (^b^ p<0.05 or ^bb^ p<0.01).

### Passive Young’s Modulus

<u>Short Lengths</u>. Analyses of passive modulus included only MHC IIA fibers. At short fiber lengths, passive modulus was significantly reduced by fatigue (19.5 ± 10.8 vs. 16.4 ± 8.3 kPa·%L_0_^-1^, respectively, p=0.006, Figure 5A). The significant interaction between fatigue and training (p=0.044) prompted assessment of the effect of training on passive modulus at short lengths in non-fatigued and fatigued fibers, separately. As a result, passive modulus was significantly higher in RT versus UT non-fatigued fibers (22.6 ± 12.3 vs. 15.1 ± 6.1 kPa·%L_0_^-1^, respectively, p=0.031) but not fatigued (p=0.406) fibers (Figure 5B). Additionally, the interaction between fatigue and biological sex (p<0.01) was statistically significant (Figure 5A), prompting post-hoc analysis via separate mixed models for males and females. These follow-up analyses (Figure 5C) revealed that passive modulus was significantly reduced by fatigue in males (23.4 ± 12.5 vs. 15.8 ± 9.2 kPa·%L_0_^-1^, respectively, p<0.001) but not females (16.2 ± 7.9 vs. 16.9 ± 7.4 kPa·%L_0_^-1^, respectively, p= 0.380). A significant fatigue x training interaction (p=0.022) in the male cohort prompted analysis of the fatigue effect in RT and UT males, separately. Passive modulus was significantly reduced by fatiguing exercise in fibers from RT (29.4 ± 13.3 vs. 19.1 ± 10.8 kPa·%L_0_^-1^, respectively, p <0.001) and UT (15.4 ± 4.2 vs. 12.3 ± 5.3 kPa·%L_0_^-1^, respectively, p = 0.024) males at short lengths.

**Figure 5:**
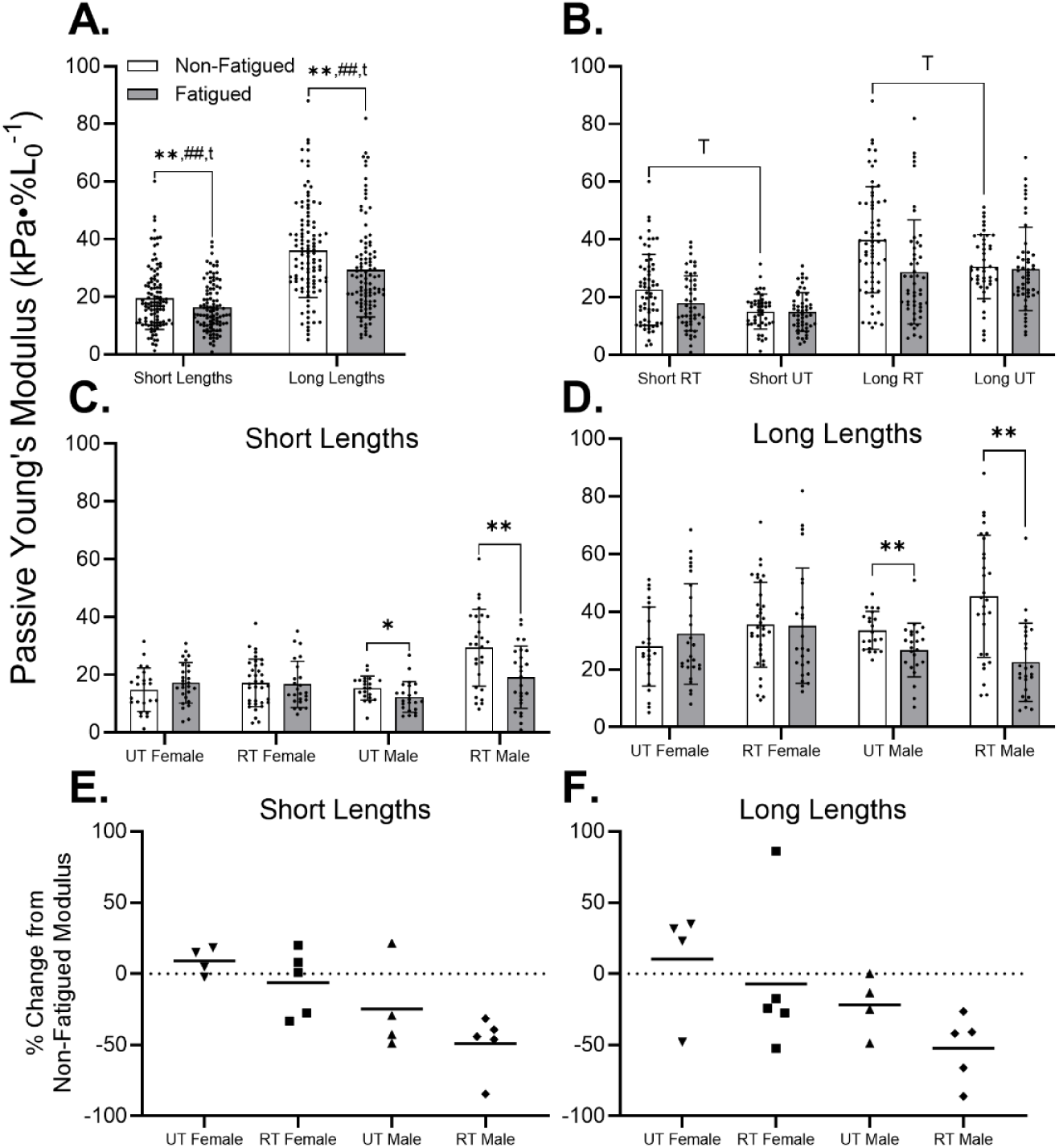
**(A)** Passive Young’s Modulus was significantly reduced by fatiguing exercise at short and long lengths. Furthermore, the biological sex by fatigue and training by fatigue interactions were significant at short and long lengths. **(B)** In non-fatigued fibers, passive Young’s Modulus was significantly greater in RT versus UT individuals at short and long lengths. At short **(C)** and long **(D)** SL, it becomes evident that reductions in passive modulus were driven by RT and UT males, but not females. On a per-individual basis, passive modulus was reduced in 9 of the 10 included males at short **(E)** and long **(F)** lengths. However, females exhibited little to no change in passive modulus at short lengths, and varied responses at long lengths. Data represent mean ± SD. Symbols indicate significant effect of fatigue (* p<0.05 or ** p<0.01), fatigue by training interaction (^t^ p<0.05), fatigue by biological sex interaction (^##^ p<0.01), or main effect of training (^T^ p<0.05).

<u>Long Lengths</u>. At long fiber lengths, passive modulus was significantly reduced by fatigue (36.1 ± 16.4 vs. 29.3 ± 16.2 kPa·%L_0_^-1^, respectively, p=0.004, Figure 5A), and was significantly altered by interactions between biological sex and condition (p<0.001) and training and condition (p=0.037). Subsequent analyses revealed that training significantly increased passive modulus of non-fatigued (39.9 ± 18.4 vs. 30.6 ± 11.1 kPa·%L_0_^-1^, respectively, p=0.045) but not fatigued (p=0.925) fibers (Figure 5B). Additionally, fatigue reduced passive modulus of fibers from males (40.3 ± 17.4 vs. 24.6 ± 11.8 kPa·%L_0_^-1^, respectively, p<0.001) but not females (p=0.296). Furthermore, the significant training x condition interaction in the male cohort revealed that passive modulus was significantly reduced by fatigue in RT (45.4 ± 21.2 vs. 22.6 ± 13.6 kPa·%L_0_^-1^, respectively, p<0.001) and UT (33.6 ± 6.5 vs. 26.8 ± 9.3 kPa·%L_0_^-1^, respectively, p=0.004) males at long lengths (Figure 5D).

Figures 4E and 4F illustrate the individual responses, expressed as a percent difference between the passive modulus of non-fatigued fibers, and that of fatigued fibers. Although RT and UT males demonstrated reduced passive cellular modulus in fatigued versus non-fatigued fibers at short (Figure 5E) and long (Figure 5F) lengths, the response in females varied considerably by individual, especially at long lengths.

### Stress Decay Index

There were no significant effects of biological sex (p= 0.657 at SL 2.6 µm, p= 0.437 at SL 2.8 µm, p= 0.968 at SL 3.0 µm, p= 0.530 at SL 3.2 µm, p= 0.193 at SL 3.4 µm, p= 0.419 at SL 3.6 µm, p=0.435 at SL 3.8 µm) or training (p= 0.452 at SL 2.6 µm, p= 0.500 at SL 2.8 µm, p= 0.752 at SL 3.0 µm, p= 0.429 at SL 3.2 µm, p= 0.269 at SL 3.4 µm, p= 0.706 at SL 3.6 µm, p=0.945 at SL 3.8 µm) on SDI at any SL. On the other hand, SDI was significantly reduced in fatigued versus non-fatigued fibers at SL 3.8 µm only (257.43 ± 171.75 vs. 303.08 ± 131.20 kPa·s·mm^-2^, p=0.034), but not at any other length (p= 0.079 at SL 2.6 µm, p= 0.656 at SL 2.8 µm, p= 0.704 at SL 3.0 µm, p= 0.151 at SL 3.2 µm, p= 0.449 at SL 3.4 µm, p= 0.074 at SL 3.6 µm, Figure 6A). Furthermore, there were significant sex by fatigue interactions at SL = 3.4 µm (p=0.021), 3.6 µm (p=0.007), and 3.8 µm (p=0.001) but not at other SLs (p= 0.871 at SL 2.6 µm, p= 0.476 at SL 2.8 µm, p= 0.188 at SL 3.0 µm, p= 0.329 at SL 3.2 µm). As a result, the effect of fatigue on SDI was examined in males and females, separately, at these SLs. In non-fatigued fibers only, there were no differences in SDI between males and females, at any SL (p= 0.804 at SL 2.6 µm, p= 0.180 at SL 2.8 µm, p= 0.424 at SL 3.0 µm, p= 0.818 at SL 3.2 µm, p= 0.640 at SL 3.4 µm, p= 0.740 at SL 3.6 µm, p=0.902 at SL 3.8 µm, Figure 6B). In males, SDI was significantly reduced in fatigued versus non-fatigued fibers at SL = 3.2 µm (96.17 ± 38.49 versus 80.13 ± 39.91 kPa·s·mm^-2^, respectively, p=0.027), SL = 3.4 µm (108.00 ± 46.54 versus 135.27 ± 50.31 kPa·s·mm^-2^, respectively, p=0.002), 3.6 µm (147.08 ± 61.32 versus 205.33 ± 75.27 kPa·s·mm^-2^, respectively, p<0.001), and 3.8 µm (195.78 ± 81.11 versus 292.59 ± 92.99 kPa·s·mm^-2^, respectively, p<0.001), but not at any other SLs (p= 0.133 at SL 2.6 µm, p= 0.257 at SL 2.8 µm, p= 0.189 at SL 3.0 µm, Figure 6C). However, fatigue had no effect on SDI at any SL in fibers from females (p= 0.326 at SL 2.6 µm, p= 0.781 at SL 2.8 µm, p= 0.469 at SL 3.0 µm, p= 0.698 at SL 3.2 µm, p= 0.394 at SL 3.4 µm, p= 0.564 at SL 3.6 µm, p=0.482 at SL 3.8 µm, Figure 6D).

**Figure 6:**
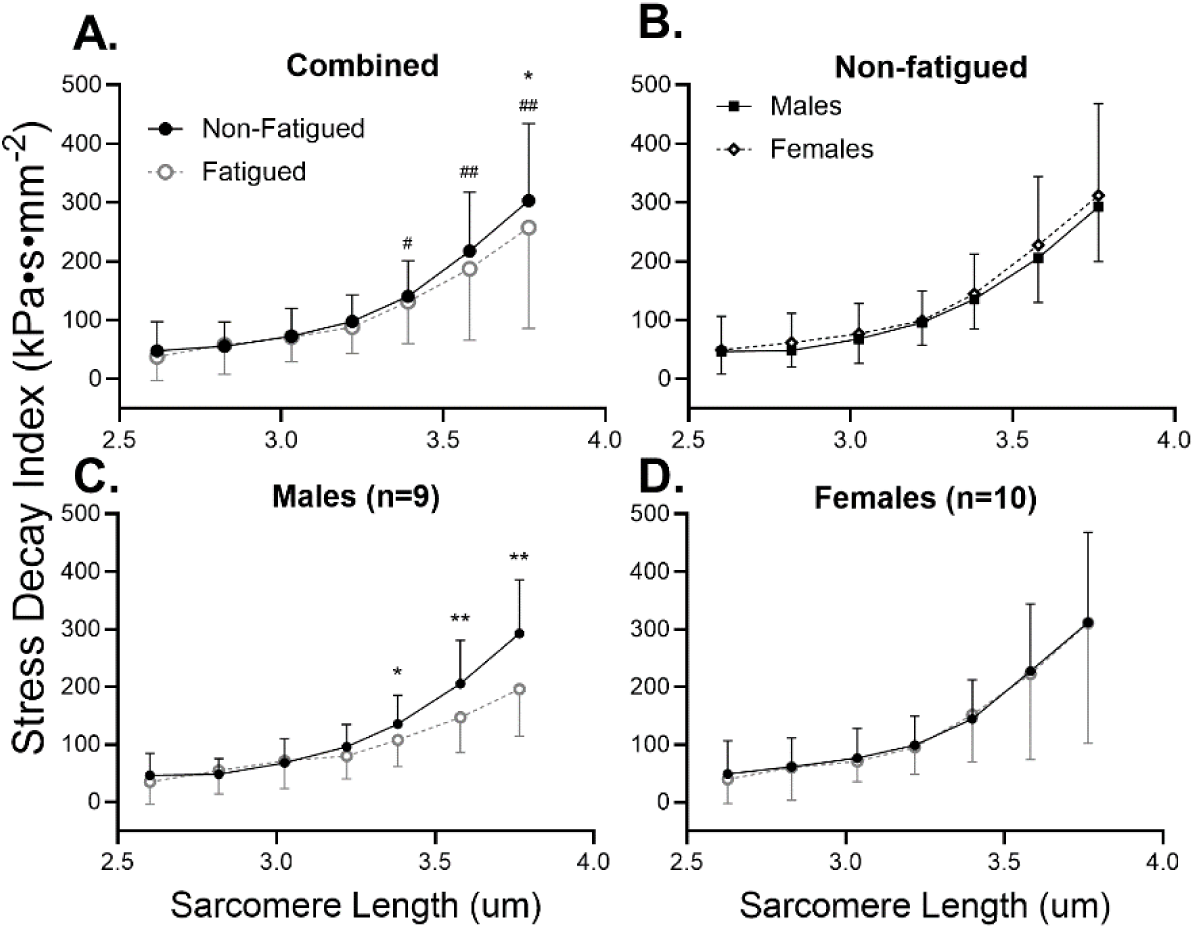
**(A)** In the combined dataset, neither biological sex nor training status had significant main effects on SDI at any SL. However, the biological sex by fatigue interaction was significant at SL=3.4 µm -3.8µm, and fatigue significantly reduced SDI at SL=3.8 µm. **(B)** In non-fatigued fibers, there was no difference in SDI between males and females, at any SL. **(C)** Stress decay index was significantly reduced by fatiguing exercise at SL=3.4 µm -3.8µm. **(D)** On the contrary, females did not exhibit changes in SDI resulting from fatiguing exercise at any SL. Data are shown as mean ± SD. Symbols indicate significant effect of fatigue (* p<0.05 or ** p<0.01) or fatigue by biological sex interaction (^#^ p<0.05 or ^##^ p<0.01).

To consider the underlying contributors to the fatigue-induced shift in SDI, we examined the measures that were used to calculate SDI: half-relaxation time, peak stress, and the magnitude of stress decay (peak stress – baseline stress). There were no significant differences in half-relaxation time due to fatiguing exercise, biological sex, or training at any SL (all p<0.05, Figure 7A). However, both peak stress and magnitude of stress decay were significantly different by fatigue, with significant interactions of biological sex by fatigue and training by fatigue. Specifically, peak stress was significantly different by fatigue (p=0.026 at SL=2.8 µm, p≤ 0.01 at SL=3.0-3.8 µm), fatigue by training interactions (p<0.01at SL 2.6, p=0.012 at SL 2.8 µm, p= 0.012 at SL 3.0 µm, p= 0.033 at SL 3.2 µm, p= 0.049 at SL 3.4 µm, p= 0.047 at SL 3.6 µm, p=0.026 at SL 3.8 µm), and fatigue by sex interactions (p=0.022 at SL=2.8 µm, p<0.01 at SL 3.0-3.8 µm, Figure 7B). Finally, the magnitude of stress decay was impacted by fatigue (p=0.030 at SL 3.4 µm, p=0.014 at SL 3.6 µm, p<0.01 at SL 3.8 µm), and the interaction of biological sex by fatigue (p=0.028 at SL 3.2 µm, p<0.01 at SL 3.4-3.8 µm) at long lengths, and by the interaction of training by fatigue at short lengths (p=0.023 at SL 2.6 µm, p=0.013 at SL 2.8 µm, p=0.042 at SL 3.0 µm, Figure 7C).

**Figure 7:**
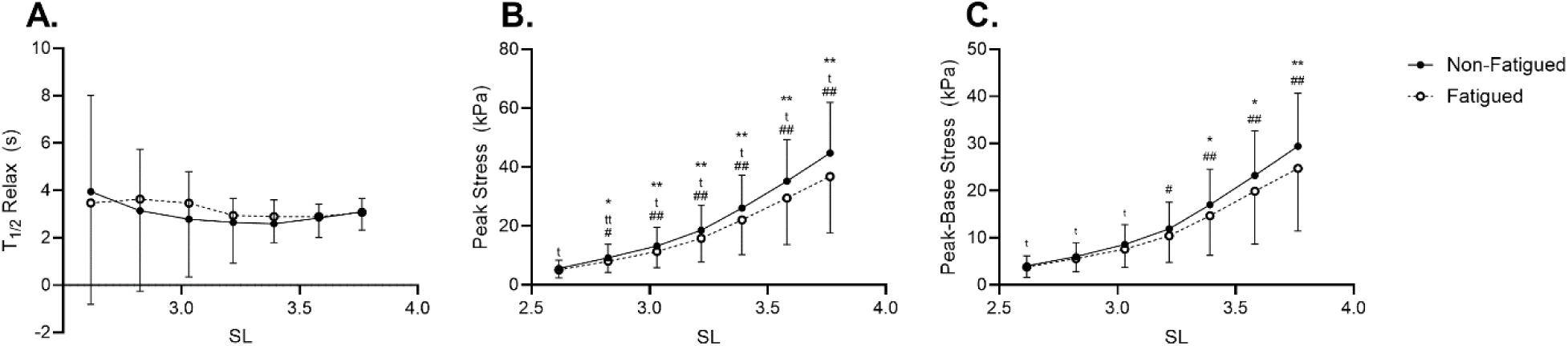
When considering the measures used to calculate SDI, it becomes clear that half-relaxation time **(A)** is not impacted by fatiguing exercise. However, both peak stress **(B)** and the magnitude of stress decay **(C)** are reduced by fatiguing exercise with both measures demonstrating mediating effects of biological sex and training status on the response to fatigue. Data are shown as mean ± SD. Symbols indicate significant effect of fatigue (* p<0.05 or ** p<0.01) fatigue by training interaction (^t^ p<0.05 or ^tt^ p<0.01), or fatigue by biological sex interaction (^#^ p<0.05 or ^##^ p<0.01).

To test whether spontaneous actomyosin interactions while in relaxing (pCa 8.0) solution contributed to observed differences in SDI, fatigue-induced differences in SDI were assessed in a subset of fibers from 2 UT young males treated with 40mM of BDM. In these BDM-treated fibers, SDI was significantly reduced in fatigued compared to non-fatigued fibers at SL = 3.4 µm (89.06 ± 19.52 versus 169.16 ± 66.95 kPa·s·mm^-2^, respectively, p=0.009), 3.6 µm (115.40 ± 35.55 versus 260.22 ± 112.20 kPa·s·mm^-2^, respectively, p=0.006), and 3.8 µm (197.54 ± 50.78 versus 425.61 ± 234.63 kPa·s·mm^-2^, respectively, p=0.026, Figure 8), consistent with observations in non-treated fibers.

**Figure 8:**
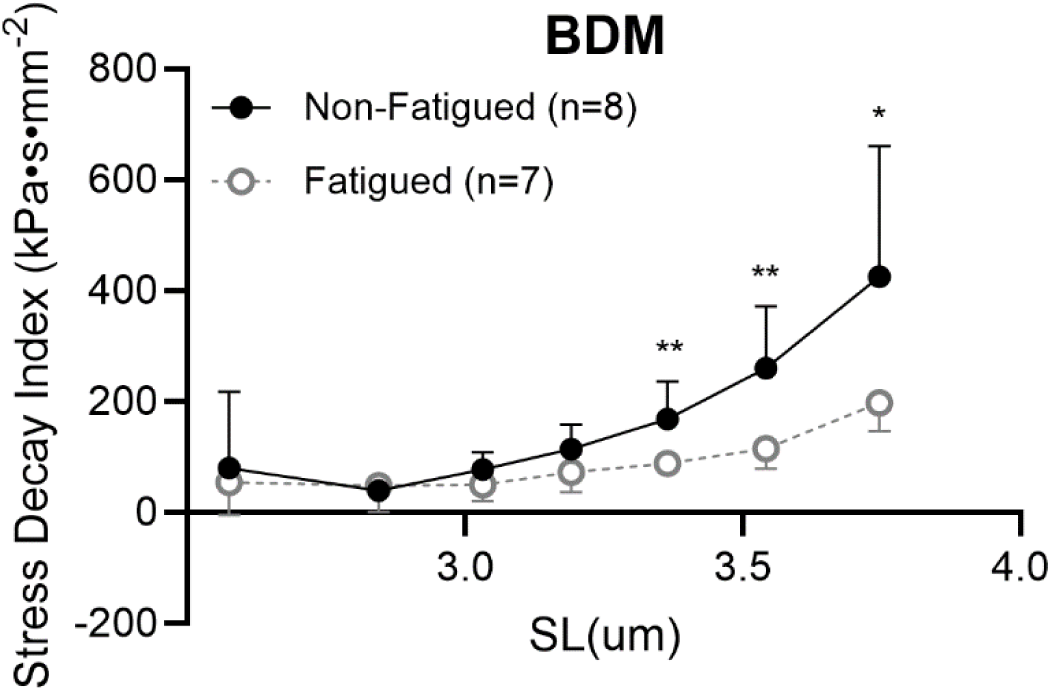
In BDM treated fibers from 2 UT males, fatigue induced differences in SDI persisted at SL 3.4-3.8 µm, suggesting that intracellular proteins other than myosin and actin contribute to this phenomenon. Data represent mean ± SD. Symbols indicate significant effect of fatigue (*, p<0.05 or **, p<0.01).

### Tissue-level Measures

Due to the linearity of the stress-strain curve for tissue samples, passive Young’s Modulus was quantified as the slope of the entire stress-strain curve. When the cohort was studied as a whole, there was no significant difference between modulus values of males and females (14.34 ± 9.67 vs. 25.91 ± 12.22 kPa·%L_0_^-1^, respectively, p = 0.197) or of non-fatigued and fatigued samples (21.06 ± 11.67 vs. 23.93 ± 13.76 kPa·%L_0_^-1^, respectively, p = 0.287). However, given the relatively limited sample in males versus females (26 vs. 58 bundles, respectively), a subsequent analysis tested the effect of fatigue condition on tissue passive modulus in females only. This sample was limited post-hoc by our desire to match bundle size across samples. In order to account for the potential effect of bundle size on passive modulus (Malakoutian *et al*., 2021), we eliminated bundles from analyses that were 2 SD below or above or below the inclusive mean for CSA (n=7). Our restricted analysis revealed a modest but significant increase in passive modulus of the fatigued versus non-fatigued bundles (28.98 ± 12.85 vs. 23.57 ± 11.36 kPa·%L_0_^-1^, p=0.036) in the female cohort but no difference in the males (NF: 18.37 ± 9.44, F: 17.28 ± 8.62 kPa·%L_0_^-1^, p=0.801, Figure 9A). Notably, the result was produced when a wider range of tissue sizes were analyzed (data not shown). To assess whether cellular changes in passive modulus resulting from fatiguing exercise were associated with the same changes in CMT modulus, average change in fiber versus CMT passive modulus was plotted for each participant for which both types of sample were tested. Although values are clustered by biological sex, and changes in fiber and CMT values with fatigue are relatively consistent, the correlation between change in fiber modulus versus that of CMT was not significant (p=0.26, Figure 9B). Finally, passive modulus measures from CMT samples of one RT female were collected before and after incubation in KI and KCl. Before treatment, passive modulus was significantly higher in fatigued versus non-fatigued CMT (30.78 ± 7.62 versus 23.59 ± 8.39 kPa·%L_0_^-1^, respectively, p=0.025, Figure 9C). However, passive modulus following KI/KCl treatment was not different between fatigued (5.17 ± 3.13 kPa·%L_0_^-1^) and non-fatigued (2.40 ± 1.88 kPa·%L_0_^-1^) CMT (p=0.370). As anticipated, KI/KCl treatment significantly and dramatically reduced passive modulus of non-fatigued and fatigued fibers (p<0.001 for both).

**Figure 9:**
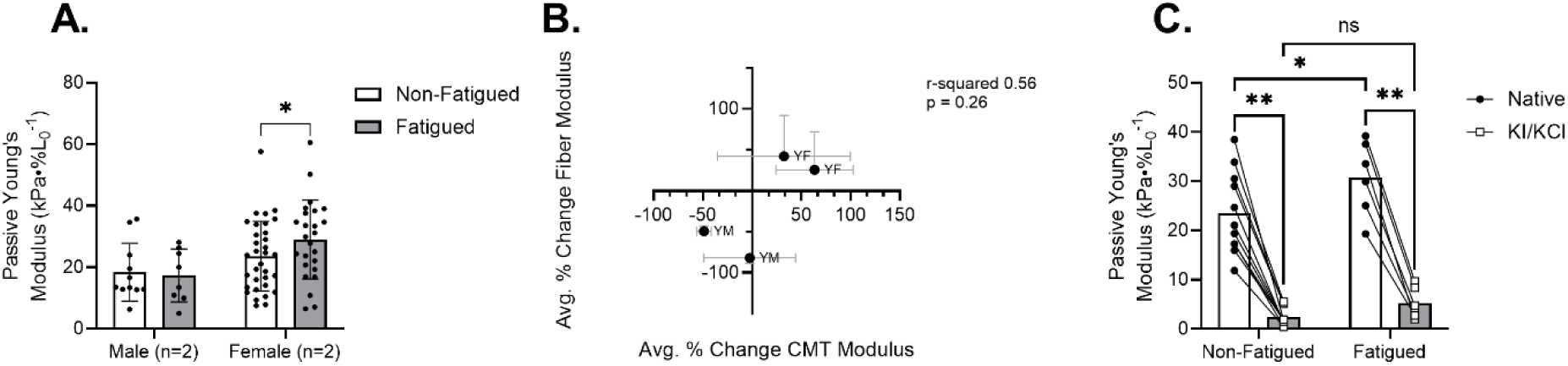
(A) Although tissue-level passive modulus was not affected by fatiguing exercise in the male cohort, it was significantly increased by fatiguing exercise in the female cohort (p=0.036). (B) The direction of change in passive modulus was similar between fibers and CMT for each participant, the correlation of change was insignificant. (B) Fatigue-induced differences in passive modulus were detected at the level of CMT, however, chemical elimination of intracellular contributors to passive modulus abolished the fatigue effect. Passive modulus was similarly reduced by treatment in fatigued and non-fatigued CMT. Each point represents one CMT sample, with lines connecting the pre-post treatment values. Symbols indicate fatigue difference (*, p<0.05 or **, p<0.001).

## Discussion

### Study sample

In the present group of participants, height, weight, and BMI were greater in males versus females (Table 1). Importantly, the step count and time spent in moderate activity per day was higher in the RT compared to UT participants (Table 1). What’s more, whole-muscle performance measures (Table 3) were higher in RT versus UT participants and in males versus females. However, there was no effect of biological sex or training on the duration of fatiguing exercise or the fatigue ratio, an index of the magnitude of fatigue (Table 3). Thus, it is unlikely that interactions between fatigue and biological sex or training are due to differences in the duration of fatiguing exercise or magnitude of in-vivo fatigue. Finally, although fiber-type distribution and fiber CSA differed across training and biological sex groups (Table 2), final analyses included only MHC IIA fibers and mechanics measures were reported normalized to CSA to minimize confounding effects on outcome measures. While restricting assessment to MHC IIA fibers somewhat limits our ability to make more broad generalizations about the applicability of our findings, we remain confident the phenomenon of fatigue-induced changes in muscle tissue mechanics reported here persists across multiple fiber types. Indeed, in separate analyses including all measured fibers, regardless of fiber type (n = 423), conclusions related to the effect of fatiguing exercise on passive stress and Young’s Modulus were not altered. However, variations by fiber type across conditions contributed significant variability that disrupted observed interactions between main effects of fatigue and training/sex. Therefore, we sought to avoid the potential for confounding interpretations, based on different fiber type distributions across groups, by limiting our assessments to samples where MHC isoform was relatively equally represented across conditions (MHC IIA, see Table 2).

### Cellular active mechanics

Active tension was not significantly different by biological sex or by training group. However, a significant fatigue by biological sex interaction revealed a modest yet significant reduction in active tension of fatigued versus non-fatigued fibers in males only (Figure 3). This has been previously observed (Privett *et al*., 2021; Ricci *et al*., 2023), and may be due to exercise induced PTM of sarcomere proteins involved in active contractile function. For example, slow skeletal myosin binding protein-C (ssMyBP-C), a thick filament regulatory protein that interacts with both the thick and thin filaments to regulate crossbridge formation (Yamamoto, 1986; Korte *et al*., 2003; Robinett *et al*., 2019) is phosphorylated by fatiguing exercise (Privett *et al*., 2021; Ricci *et al*., 2023). Whereas dephosphorylated ssMyBP-C is more likely to bind to both the thick and thin filaments, resulting in slowed crossbridge kinetics, phosphorylation releases this clutch-like mechanism, enhancing crossbridge kinetics (Colson, 2019; Robinett *et al*., 2019). Therefore, it is plausible that accelerated crossbridge kinetics resulting from ssMyBP-C phosphorylation during fatiguing exercise may reduce force production. Though we can only speculate as to the mechanisms explaining reduced isometric tension of fatigued fibers in male participants, tension production in these fibers clearly demonstrates adequate function of all fibers included in the present analyses.

### Chronic Training Increases Baseline Passive Stress and Young’s Modulus at Longer Lengths

In non-fatigued fibers, passive stress (Figure 4B) and Young’s modulus (Figure 5B) were higher in fibers from RT versus UT participants at longer SLs. This observation is consistent with previous work (Noonan *et al*., 2020), in which training was shown to modify cellular passive modulus in rodent skeletal muscle fibers in a training-modality-dependent manner. While chronic training may impact ECM-based stiffness in skeletal muscle (Kjær, 2004), this is unlikely to influence the cellular mechanics reported in this present study due to the use of chemically permeabilized skeletal muscle fibers, which no longer have intact ECM. Rather, it is more likely that intracellular proteins are responsible for the observed training effect on passive measures. The fact that training effects were observed at longer SLs (3.0-3.8µm), where overlap of thick and thin filaments is minimal, suggests that the sarcomere protein titin is at least partially involved.

Differential expression of titin isoforms across participants have previously been reported (Fry *et al*., 1997), presenting the possibility that training-induced shifts in titin isoform distribution may have contributed to the observed increase in passive stress or modulus in RT versus UT participants. Yet, existing literature does not support the notion of training-induced shifts in titin isoform (McGuigan *et al*., 2003; Pellegrino *et al*., 2016) or titin size (Noonan *et al*., 2020) in skeletal muscle. Nonetheless, 3 weeks of exercise training has been demonstrated to shift titin isoform in murine cardiac muscle (Hidalgo *et al*., 2014) suggesting that the effect of exercise on titin isoform may be specific to training modality and muscle type. It is also possible that chronic training alters titin-based stiffness through changes to titin PTMs. Training-based differences in titin phosphorylation, a well-studied mediator of titin-based stiffness, have been observed in murine cardiac muscle (Hidalgo *et al*., 2014). However, it is not yet known whether chronic training alters the phosphorylation background of titin in human skeletal muscle, and we did not report titin phosphorylation measures in the present study. Although it is unclear how chronic training impacts skeletal muscle titin in humans, specifically, there is certainly a role for titin in skeletal muscle mechano-signaling and remodeling in response to exercise (Krüger & Kötter, 2016), supporting the notion that titin-based stiffness may adapt to a chronic training stimulus in order to better meet the demands of intense physical exercise.

### Fatiguing Exercise Alters Cellular Passive Mechanics

In support of our initial hypothesis, fatiguing exercise significantly reduced both passive stress (Figure 4A) and Young’s Modulus (Figure 5A) when all participants were considered collectively. Given that the effect of fatiguing exercise on passive stress and modulus was mediated by biological sex and training status, subsequent analyses examined training- and biological sex-based groups separately. As a result, it became clear that trained males had the most robust reduction in passive stress (Figure 4F) and modulus (Figure 5C-D) in fatigued versus non-fatigued fibers, yet untrained females exhibited a non-significant upward trend in passive stress (Figure 4C) and modulus (Figure 5D) in fatigued versus non-fatigued fibers. In all groups where a mean difference (even a non-significant one) in passive stress or modulus was observed between non-fatigued and fatigued fibers, the difference increased as SL increased. In fact, the difference in mean passive stress between fatigued and non-fatigued fibers was largest at longer SLs (∼3.4 µm and longer), where thick and thin filaments no longer overlap. Although the contribution of residual actomyosin interactions to cellular passive mechanical properties is possible, we have previously demonstrated that cellular passive modulus is still altered by fatiguing exercise in fibers treated with 2,3-butanedione monoxime (BDM), a myosin inhibitor, suggesting that residual crossbridge formation is not a primary mechanism of fatigue-induced changes to cellular passive mechanics (Privett *et al*., 2024). Rather, the fact that fatigue-induced differences are largest at longest lengths, where passive mechanics in permeabilized muscle fibers are predominantly titin-based, suggests that titin-based mechanisms may have contributed to these observations. Titin-based stiffness can be acutely regulated via PTM, and it is possible that exercise-induced small molecules such as inorganic phosphate (Müller *et al*., 2014; Hamdani *et al*., 2017), oxidative molecules (Alegre-Cebollada *et al*., 2014), or heat shock proteins (Kötter *et al*., 2014) bind to viscoelastic regions of titin during fatiguing exercise, thereby altering titin-based viscoelasticity.

We have previously reported altered titin phosphorylation following a single bout of fatiguing exercise (Privett *et al*., 2024), which may have implications for titin-based stiffness (Hamdani *et al*., 2017). However, the specific response of titin-based stiffness to phosphorylation is highly dependent on the location at which the phosphorylation event occurs, as has been observed previously (Müller *et al*., 2014). Separately, S-glutathionylation of cryptic binding sites of cardiac titin has been shown to inhibit titin Ig domain refolding, thereby reducing titin-based stiffness (Alegre-Cebollada *et al*., 2014). In skeletal muscle, S-glutathionylation has been demonstrated to reduce passive force resulting from fiber stretch, yet the observed reductions were smaller than those observed in cardiac muscle (Watanabe *et al*., 2020). However, oxidation events such as S-glutathionylation are not likely present in our sample due to the use of the antioxidant DTT in dissecting and relaxing solutions. Finally, it is possible that small HSPs generated during fatiguing exercise may bind to titin in a way that alters titin-based stiffness. Previous work reported the translocation of small HSPs, alphaβ-crystallin and HSP27, to the elastic regions of skeletal muscle titin (36). In the same study, the binding of these small HSPs in cardiac myocytes prevented acidification-based aggregation of titin Ig domains and subsequent increases in titin-based stiffness. Our present study did not measure changes to titin PTMs, precluding the ability to draw conclusions regarding their contribution to the observed changes in passive stress and modulus.

However, these passive measures were made in permeabilized single skeletal muscle fibers, suggesting the observed differences were at least in part due to a titin-based mechanism. Further studies in this area should seek to determine where exercise-induced changes to titin phosphorylation occur along the titin protein (Privett *et al*., 2024) and should assess whether the binding of small HSPs to the elastic regions of skeletal muscle titin is altered by an acute bout of fatiguing exercise.

*Biological Sex and Chronic Training Mediate the Effect of Fatiguing Exercise on Cellular Passive Stress and Young’s Modulus.* Neither the duration of fatiguing exercise nor the fatigue ratio differed by biological sex or training status; therefore, the sex- and training-based differences in response to fatiguing exercise are not likely explained by differences in performance of the fatiguing exercise itself (Table 3).

Rather, the lack of a significant mean response in females was at least partially due to the variability among female participants (Figure 5E), especially at long lengths (Figure 5F). One potential explanation for this varied response might be sex-based differences in circulating levels of estrogen (Bell *et al*., 2011, 2012; Ham *et al*., 2020). Specifically, despite our efforts to minimize variability by collecting biopsy samples from the same point of the menstrual cycle (pre-follicular phase, n=5) or from participants using hormonal contraception (n=5), inter-individual differences in systemic estrogen may have contributed to differences in skeletal muscle mitochondrial function (Pellegrino *et al*., 2022; Yoh *et al*., 2023).

Mitochondrial function can affect the concentration of heat shock proteins (Liu & Steinacker, 2001) and oxidative molecules such as glutathione (Marí *et al*., 2009), both of which have the capacity to alter titin-based stiffness via PTM, in the sarcoplasm. Given the potential impact of varied blood estrogen concentration on intracellular mediators of skeletal muscle stiffness, future studies will seek to measure circulating estrogen concentration at the time of muscle biopsy.

The response of cellular passive mechanics was also mediated by training status. Although training-induced changes to skeletal muscle titin isoform are possible, there is no evidence for training-induced shifts in skeletal muscle titin in the literature (McGuigan *et al*., 2003; Pellegrino *et al*., 2016). However, training may modify the pattern of PTMs to titin. There is evidence that titin-based stiffness can be modified by phosphorylation (Hamdani *et al*., 2017) and S-glutathionylation (Alegre-Cebollada *et al*., 2014), yet there is no direct evidence suggesting that chronic training results in baseline changes to either. HSPs, on the other hand, have been demonstrated to increase in baseline expression following training (Morton *et al*., 2009). Therefore, it is possible that HSPs exert a greater influence in the skeletal muscle of trained versus untrained young adults. For this reason, future studies should seek to measure HSP binding to titin in human VL samples.

### Stress Decay Index

Although the incremental stretch protocol used here precludes the measurement of viscosity, per se, SDI is a useful proxy for assessing the impact of fatiguing exercise on VL viscoelasticity (Lim *et al*., 2019). In the results reported here, there is a clear mediating effect of biological sex on the response of SDI to fatiguing exercise such that fatigue reduces SDI at longer SL in samples from males but not females (Figure 6). To interrogate the underlying contributors to fatigue-induced alteration to SDI, half-relaxation time, peak stress, and magnitude of stress decay were analyzed (Figure 7). There was no effect of fatigue, biological sex, or training on half-relaxation time, suggesting that these factors did not impact the time course of stress decay. Rather, the peak stress and the magnitude of stress decay were both reduced in fatigued versus non-fatigued fibers, with mediating effects of training and/or biological sex on fatigue response in a length-dependent manner. In skeletal muscle myofibrils, viscoelastic behavior arises from titin immunoglobulin domain unfolding such that the amplitude of force decay increases as sarcomere length increases (Minajeva *et al*., 2001), paralleling the results presented here (Figure 7C). The possibility that residual actomyosin binding interactions in our relaxing solution (pCa 8.0) may have contributed to the observed differences in SDI was considered. However, the effect of fatigue on SDI was still present in BDM-treated fibers (Figure 8), suggesting that actomyosin binding was unlikely to be a primary contributor to fatigue-induced differences in SDI. The present study design does not allow us to draw strong conclusions regarding the physiological importance of fatigue-induced changes to titin viscoelasticity to whole-muscle function. However, one previous study utilized a unique knock-out model targeting the PEVK region, a prominent contributor to cardiac viscoelasticity, to produce a 40% reduction in left ventricular chamber viscosity in the PEVK knock-out hearts (Chung *et al*., 2011). Although it is important to acknowledge the potential muscle specificity of the mechanisms and physiological relevance of titin-based viscoelastic properties, this work highlights the potential role of titin in modulating whole-muscle viscoelasticity. As such, future studies should include methods to measure true viscosity of fatigued and non-fatigued skeletal muscle cells and composite tissue.

### Contribution of intracellular proteins to tissue-level passive modulus

To explore how altered cellular passive modulus might impact mechanics at the tissue level, the effect of fatiguing exercise on CMT samples was considered in a subset of males and females (Figure 9A). e, CMT modulus was not different following fatigue in males but was elevated following fatigue in females. Though seemingly in contrast to fiber level data in the more complete sample, this observation was consistent with fiber level differences from the four individuals included in this subset. This parallel response in fiber and CMT modulus to fatiguing exercise supported our interpretation that intracellular mediators of passive modulus contribute to passive mechanics at higher levels of organization, though we were unable to directly assess their relative contribution from the data presented here (Figure 9B). These associations were not significant and are interpreted with some caution, given prior evidence that the inclusion of ECM (Mathewson *et al*., 2014; Ward *et al*., 2020) may obscure the influences of the myofiber. CMT samples from one RT female were subjected to passive stretch before and after treatment with KI and KCl to eliminate sarcomere thin and thick filaments, respectively. The removal of thick and thin filaments via KI/KCl is an established method (Ottenheijm *et al*., 2012) to eliminate titin-based contributions to cardiac and skeletal muscle stiffness. Therefore, any remaining passive force measured following treatment is attributed to ECM. In the subset of CMT samples used for KI/KCl experiments, CMT passive modulus was higher in fatigued versus non-fatigued samples before treatment (Figure 9C), similar to observations in single fibers from this same individual. Following KI/KCl treatment, fatigue-induced differences in modulus were no longer evident, suggesting that differences in fatigued versus non-fatigued passive modulus in CMT samples were dependent on intracellular mechanisms including titin. Indeed, passive modulus following KI/KCl was dramatically reduced in all samples, highlighting the importance of intracellular mechanisms to CMT modulus in these bundles of 12-14 fibers. Though these exploratory measures with reduced sample size limit definitive conclusions in this study, previous studies of genetically modified murine muscle tissue have shown direct effects of titin on whole-muscle passive tension (Brynnel *et al*., 2018), supporting the idea that alterations in titin mechanics may scale to the whole tissue level. Further study is required to determine how these changes manifest when scaling from in-vitro cellular measures to in-vivo tissue levels of organization (Ward *et al*., 2020). With those caveats in mind, the data in the present manuscript contribute to a growing body of literature supporting the notion that intracellular mechanisms contribute to whole muscle tissue mechanics in vivo, encouraging further research into the potential magnitude and dynamic regulation of these effects.

## Limitations

The present findings support earlier data (Privett *et al*., 2024) suggesting a sex-specific response to acute muscle fatigue that reduces viscoelasticity in males, but not females. These acute responses are observed regardless of the presence or absence of chronic effectors impacting viscoelasticity (physical training history; Figure 4B). Sex based differences in the dynamic response to acute physiological stressors is notable, but caution is warranted when attempting to extrapolate these findings to in-vivo observations. Though we examined elastic modulus in CMT samples from a subset of volunteers and demonstrated the important role of fibrillar mechanics to tissue level behaviors (Figure 9C), we have not replicated our primary finding in CMT. Therefore, further experiments are required to explore how altered titin-based mechanics might impact muscle tissue that includes ECM and associated structures in-vivo. Further, additional experiments are needed to clarify whether passive mechanics in permeabilized single fibers might reflect in-vivo function that may impact injury prevention or performance. Indeed, while a growing body of literature supports the notion that titin plays a critical role in stretch-shortening dynamics at the fiber level (Nishikawa *et al*., 2013; Power *et al*., 2013; Herzog, 2014; Herzog *et al*., 2016; Nishikawa, 2016, 2020), it is not clear at present whether alterations observed here, under passive conditions, may impact performance during dynamic loading.

## Conclusions

In conclusion, fatiguing exercise impacts cellular passive stress and Young’s Modulus in a way that is mediated by training status and biological sex in young adults. Whereas trained and untrained young males demonstrated reduced passive stress and Young’s Modulus in fatigued versus non-fatigued single fibers, neither trained nor untrained young females demonstrated a change in mean stress or Young’s Modulus at any length. However, a look at individual differences in mean Young’s Modulus of fatigued versus non-fatigued fibers revealed that females exhibited a high degree of variability in the direction and magnitude of change in Young’s Modulus. We speculate that inter-individual differences in circulating estrogen may have contributed to this variability; however, future studies will need to measure blood estrogen at time of study to support or refute this idea. Like passive stress and modulus, the effect of fatiguing exercise on cellular SDI was mediated by biological sex, with only males demonstrating significantly reduced SDI in fatigued versus non-fatigued fibers at SLs 3.4-3.8 µm due to changes in the magnitude of stress decay, rather than changes to the time course of stress decay. Finally, CMT assays suggest that although intracellular proteins appear to impact CMT passive modulus, their relative impact on the effect of fatiguing exercise on CMT passive mechanics is not yet clear.

## Data availability statement

Supporting data in results are presented in the manuscript and included in figures. Additional data are available as supporting information published with the paper online.

## Acknowledgements

The Authors wish to thank the volunteers for their invaluable contributions to our work. Additionally, thanks are due to Jordan Cooper, who provided valuable assistance in experimental assays and data analysis associated with the present study.

## Author Contributions

These experiments were conducted in the Muscle Cellular Biology Laboratory in the University of Oregon Human Physiology Department. G.E.P., A.W.R., K.W.N, and D.M.C. designed and performed experiments. G.E.P., A.W.R., K.W.N, and D.M.C. analyzed and interpreted study data and revised the manuscript. G.E.P. drafted the manuscript. D.M.C. conceived of and directed the study. All authors approved the final version of the manuscript and agree to be accountable for all aspects of the work in ensuring that questions related to the accuracy or integrity of any part of the work are appropriately investigated and resolved. All persons designated as authors qualify for authorship and all those who qualify for authorship are listed.

